# Reconstruction and deconstruction of human somitogenesis in vitro

**DOI:** 10.1101/2022.05.11.491561

**Authors:** Yuchuan Miao, Yannis Djeffal, Alessandro De Simone, Kongju Zhu, Andrew Silberfeld, Jong Gwan Lee, Jyoti Rao, Oscar A. Tarazona, Alessandro Mongera, Pietro Rigoni, Margarete Diaz-Cuadros, Laura Min Sook Song, Stefano Di Talia, Olivier Pourquié

**Affiliations:** Department of Genetics, Harvard Medical School and Department of Pathology, Brigham and Women’s Hospital, Boston, MA, USA; Department of Cell Biology, Duke University Medical Center, Durham, NC, USA; Harvard Stem Cell Institute, Harvard University, Cambridge, MA USA

## Abstract

The body of vertebrates displays a segmental organization which is most conspicuous in the periodic organization of the vertebral column and peripheral nerves. This metameric organization is first implemented when somites, which contain the precursors of skeletal muscles and vertebrae, are rhythmically generated from the presomitic mesoderm (PSM). Somites then become subdivided into anterior and posterior compartments essential for vertebral formation and segmental patterning of the peripheral nervous system^1–4^. How this key somitic subdivision is established remains poorly understood. Here we introduce novel tridimensional culture systems of human pluripotent stem cells (PSCs), called Somitoids and Segmentoids, which can recapitulate the formation of epithelial somite-like structures with antero-posterior (AP) identity. Using these systems, we identified a key organizing function of the segmentation clock in converting temporal rhythmicity into the spatial regularity of anterior and posterior somitic compartments. We show that an initial salt-and-pepper expression pattern of the segmentation gene *MESP2* in the newly formed segment is transformed into defined compartments of anterior and posterior identity via an active cell sorting mechanism. Moreover, we demonstrate a large degree of independence of the various patterning modules involved in somitogenesis including the segmentation clock, somite epithelialization and AP polarity patterning. Together we put forward a novel framework accounting for the symmetry breaking process initiating somite polarity patterning. Our work provides a valuable platform to decode general principles of somitogenesis and advance knowledge of human development.

---

Our peripheral nerves exhibit a striking periodic organization which coincides with that of vertebrae. This arrangement can be traced back to the original body segmentation resulting from somite formation. Somites, which form from the presomitic mesoderm (PSM), define the prepattern on which vertebral metamery is established^1^. They are repeatedly arrayed in two bilaterally symmetric columns which give rise to the skeletal muscles and axial skeleton. The PSM, which is initially mesenchymal in the posterior part of the embryo, becomes progressively epithelial as it matures. At its anterior tip, somites form rhythmically as epithelial blocks surrounding a mesenchymal core. The periodicity of somite formation involves a molecular oscillator called the segmentation clock^1,5^. This oscillator controls the rhythmic activation of Notch, Wnt and FGF pathways which manifest as traveling waves of target gene expression in the posterior PSM. These periodic signals are interpreted at the level of the determination front, whose position is defined by posterior gradients of FGF and Wnt signaling in the PSM. This “Clock and Wavefront” mechanism eventually leads to the activation of the transcription factor MESP2 in a stripe which prefigures the future segment. MESP2 is next involved in the subdivision of forming somites into an anterior and posterior compartment^6,7^. This partition is critical for peripheral nervous system segmentation as the migration of neural crest cells and peripheral axons is initially restricted to the anterior somitic compartment^8^. It is also essential for vertebrae which form from the fusion of a posterior somite compartment with the anterior compartment of the next posterior somite^3^. The mechanism controlling the formation of these anterior and posterior somitic domains remains poorly understood.

Our understanding of vertebrate segmentation only relies on studies performed in model organisms such as mouse, chicken, and zebrafish embryos. Very little is known about human somitogenesis which takes place very early during pregnancy, between 3- and 4-weeks post-fertilization^9^. The recent development of in vitro systems recapitulating paraxial mesoderm development from pluripotent stem cells (PSCs) demonstrated a high degree of conservation of the gene regulatory networks involved in PSM patterning between mouse and human embryos^10–15^. Monolayers of human PSCs differentiating to a PSM fate in vitro recapitulate the posterior FGF and Wnt gradients and the oscillations of the segmentation clock with a ∼5h period. However, a limitation of these 2D systems is that they do not allow examination of the morphogenesis of the tissues generated in vitro. In mouse, a striking recapitulation of all somitogenesis stages including epithelial somite and antero-posterior (AP) compartments formation has been achieved in 3D organoids that contain cells of all three germ layers^16,17^. However, no such protocols have so far been reported for human PSCs. Thus, whether the mechanisms involved in somite formation and patterning described in embryos of other vertebrates are conserved in humans remains unknown.

To study human somitogenesis, we set out to develop PSC-derived 3D culture systems. We first generated human iPSC spheroids in suspension, and then treated them with the Wnt agonist CHIR and the BMP inhibitor LDN for 48 hours to induce the PSM fate (Fig.1a). Subsequently we transferred the spheroids to a laminin coated substrate and used confocal microscopy to characterize gene expression dynamics as the spheroids spread out (Extended Data Fig.1a, Supplementary Video1). We used an iPS cell line harboring a destabilized Achilles (YFP) reporter at the *HES7* locus to detect segmentation clock oscillations and a mCherry reporter at the *MESP2* locus to monitor the onset of segmental determination^10^ (Extended Data Fig.1b). Live imaging showed that *HES7*, a core component of the segmentation clock, starts to oscillate with a 4-5 hour period (Fig.1b,c, Extended Data Fig.1c) as the spheroids spread out. *HES7* signals initiated from the peripheral region of the spreading organoid and propagated as concentric waves toward the center (Fig.1b, Extended Data Fig.1c, Supplementary Video2). After about 4 cycles of *HES7* oscillations, between ∼ 64h and 72h, expression of the reporter ceased and the *MESP2* reporter became simultaneously expressed across spheroids (Fig.1b,c, Supplementary Video2). Thus, the onset of *MESP2* immediately follows the arrest of *HES7* oscillations. *PAX3* is a transcription factor first expressed in the anterior PSM and epithelial somites soon after activation of *MESP2*^18^. In the differentiating organoids of a *PAX3-YFP* reporter line, we observed the onset of *PAX3* activation in all cells around 78h (Fig.1c). At 90h, numerous *PAX3*-positive somite-like epithelial rosettes started to emerge and they were visible under bright-field microscopy by 120h (Fig.1d,e). These rosettes displayed typical somitic features such as enriched apical N-Cadherin and F-actin, a laminin-rich basal lamina, and a core region filled with mesenchymal cells (Fig.1e-g, Extended Data Fig.1d,e). We performed RNAseq at 48h, 66h, and 120h of the differentiation protocol, and observed the expression of signature genes associated with PSM, Determination Front, and somites respectively (Fig.1h and Extended Data Fig.1f). Therefore, these organoids, which we term “Somitoids”, successfully recapitulate the timely progression of gene expression from PSM to somites as well as major aspects of epithelial somite morphogenesis.

**Fig. 1.**
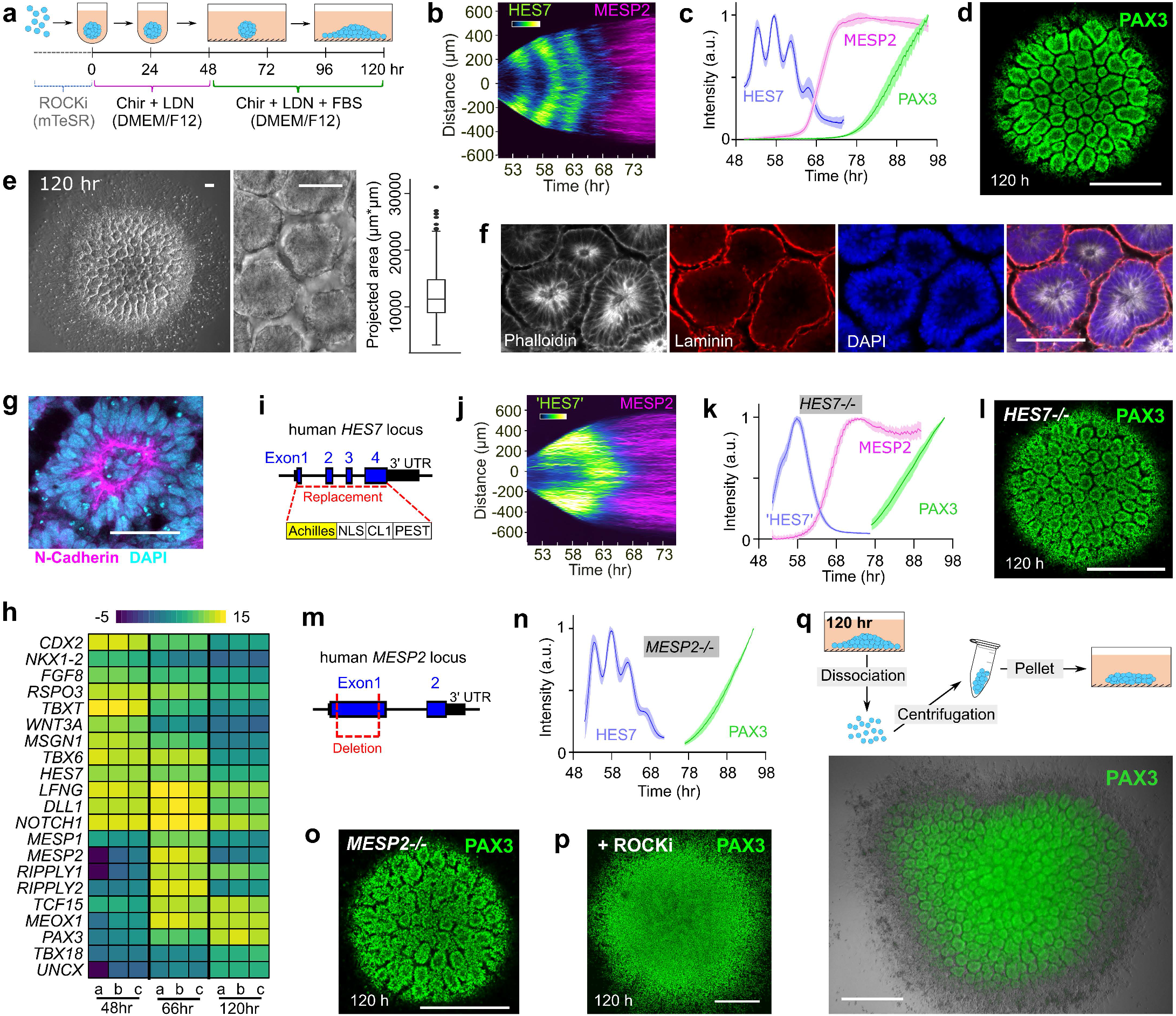
Characterization of the Somitoid model. **a**, Illustration of the Somitoid protocol. **b**, Kymograph of HES7 and MESP2 reporters obtained from a line scan across the center of a Somitoid. **c**, Temporal profiles (mean±s.d.) ofreporters for HES7 (n=5 Somitoids), MESP2 (n=6 Somitoids), and PAX3 (n=6 Somitoids). **d**, Confocal image of a PAX3-reporting Somitoid at 120h. **e**, Bright field images of 120h Somitoid and box plot of rosette projected areas (n=310 rosettes from 5 Somitoids). **f-g**, Confocal images of immunostaining of 120h rosettes. **h**, Heat map of selected genes associated with somitogenesis in 48h, 66h, and 120h Somitoids (48 Somitoids in each time point; n=3 independent experiments), as measured by RNA sequencing. Expression levels were calculated by log2 (TPM+0.01). **i**, Strategy for making HES7 knockout line. **j**, Kymograph of pseudoHES7 and MESP2 reporters in a HES7-null Somitoid. **k**, Temporal profiles (mean±s.d.) of reporters for pseudoHES7 (n=6 Somitoids), MESP2 (n=6 Somitoids), and PAX3 (n=9 Somitoids) in HES7-null Somitoids. **l**, Confocal image of a PAX3-reporting HES7-null Somitoid at 120h. **m**, Strategy for making MESP2 knockout line. **n**, Temporal profiles (mean±s.d.) of reporters for HES7 (n=6 Somitoids) and PAX3 (n=8 Somitoids) in MESP2-null Somitoids. **o**, Confocal image of a PAX3-reporting MESP2-null Somitoid at 120h. **p**, Confocal image of a PAX3-reporting WT Somitoid treated with 10μM ROCKi. **q**, Experiment scheme (top) and wide-field image (bottom) of re-aggregating 120h Somitoids. An overlay image of PAX3 reporter fluorescence and bright field is shown. Scale bars represent 500μm (d, l, o, p, q), 100μm (e, f), and 50μm (g).

To explore the role of the segmentation clock in rosette formation, we replaced the coding sequence of *HES7* with a destabilized Achilles (YFP) reporter to generate a null mutant (Fig.1i). The YFP signal thus represents the activity of the *HES7* promoter in absence of HES7 protein, and we confirmed that the periodic dynamics was ablated (Fig.1j,k, Supplementary Video3). Yet the *HES7*-null explants proceeded with sequential expression of *MESP2* and *PAX3* followed by rosette formation as observed in controls (Fig.1j-l). We next generated a *MESP2-*null mutant iPS line which exhibited normal *PAX3* expression and generated epithelial rosettes similar to wild type controls (Fig.1m-o). Formation of the rosettes could be blocked without altering *PAX3* expression by inhibiting myosin contractility using Y-27632 (ROCKi) or Blebbistatin (Fig.1p, Extended Data Fig.1g). We also dissociated the Somitoids to single cells after rosettes appeared, and then re-aggregated cells by centrifugation prior to culture (Fig.1q). Strikingly, cells re-formed similar rosettes in the new aggregates (Fig.1q). Together, these experiments suggest that rosette formation is an acto-myosin dependent self-organizing property of cells differentiated to the somite stage and does not depend on a prior prepattern established by the clock and wavefront system.

We next investigated whether the epithelial rosettes exhibit an AP polarity as observed in somites. We used an iPS line harboring a *MESP2* reporter (mCherry) to mark the nascent anterior compartment and a *UNCX* reporter (YFP) for mature posterior identity (Extended Data Fig.2a). At 120h, we observed rosettes mostly composed of either YFP-high/mCherry-low or mCherry-high cells (Fig.2a). As reported in mouse embryos, *UNCX* trailed *MESP2* expression in time (Fig.2b, Supplementary Video4). RNAseq performed on FACS-sorted YFP-high and mCherry-high cell fractions from 120h cultures showed that they express signature genes associated with posterior (*UNCX, DLL1*) and anterior somite (*FGFR1, TBX18*) compartments (Fig.2c, Supplementary Information). In mouse, Notch signaling is required for *Mesp2* expression and the establishment of AP identities^7,19^. Accordingly, treatment of cultures with the Notch inhibitor DAPT prevented expression of *UNCX* and *MESP2* and rosette formation but not *PAX3* expression (Extended Data Fig.2b,c). In the presence of ROCKi or Blebbistatin, no rosette formed but YFP and mCherry-positive cells still appeared and aggregated into separate clusters (Fig.2d, Extended Data Fig.2d). *HES7*-null Somitoids showed similar *UNCX* expression and patterning as WT (Fig.2e, Extended Data Fig.2e). Further, *MESP2* deletion resulted in an expansion of *UNCX* positive cells and formed only rosettes exhibiting a posterior identity (Fig.2e, Extended Data Fig.2e), consistent with the reported role of *MESP2* in inhibiting the posterior fate to promote the anterior one^19^. Therefore, human iPS cells differentiating to the somitic fate in this in vitro system acquire distinct AP identities. However, unlike embryos, these identities do not coexist within the same epithelial somite but are mostly found in distinct epithelial rosettes. These experiments argue that acquisition of the anterior and posterior fates operates independently of the segmentation clock and rosette morphogenesis.

**Fig. 2.**
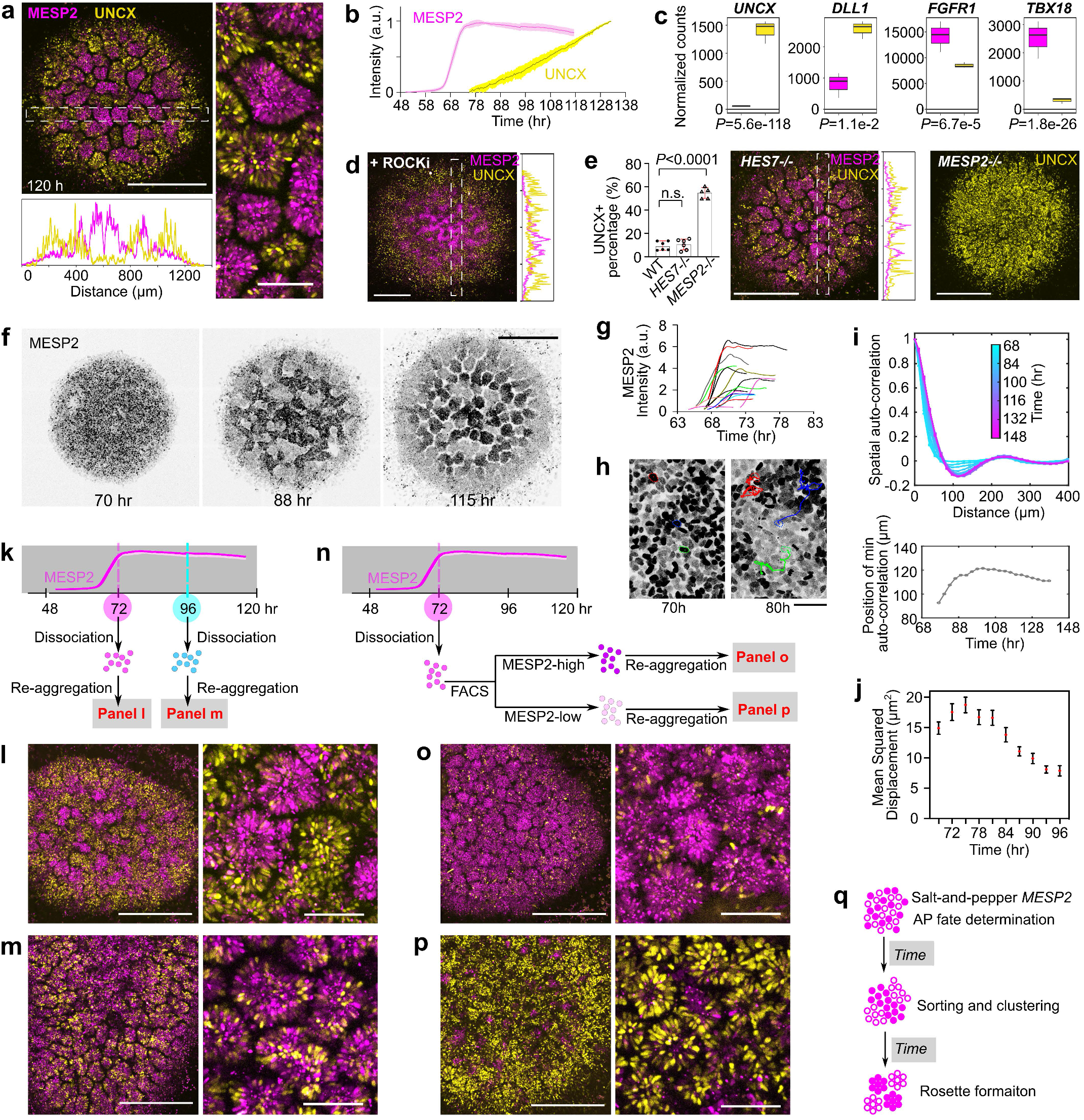
Antero-Posterior polarity patterning in the Somitoid model. **a**, Maximum z-projection images of reporters for MESP2 (magenta) and UNCX (yellow) in 120h Somitoids, and intensity profiles of MESP2 and UNCX across the dotted line box. **b**, Temporal profiles (mean±s.d.) of reporters for MESP2 (n=6 Somitoids) and UNCX (n=10 Somitoids) over the entire Somitoids. **c**, Normalized RNA counts of selected polarity genes in cell fractions separated by flow cytometry, as measured by RNA sequencing (n=3 independent experiments, 96 Somitoids in each n). Cells with top 10% mCherry fluorescence are shown on the left (magenta) and top 10% YFP fluorescence on the right (yellow). All four genes were identified as differentially expressed genes by DESeq2 using the Wald test. **d**, Image and Intensity plots of MESP2/UNCX reporters in a Somitoid treated with 10μM ROCKi. **e**, Left, percentage (mean±s.d.) of UNCX-positive cells characterized by flow cytometry in 120h WT (n=6 experiments), HES7-null (n=6 experiments), and MESP2-null (n=5 experiments) Somitoids, one-way ANOVA. Right, images of MESP2 and UNCX reporters in HES7-null Somitoids, and UNCX reporter in MESP2-null Somitoids. **f**, Time-lapse images of MESP2 reporter in a Somitoid. **g**, Temporal profiles of MESP2 reporter in individual cells. **h**, Images of MESP2 reporter in the same region. Tracks of MESP2-high cells are imposed on the 80h image with dotted outlines indicating cell positions at 70h. **i**, Top, spatial auto-correlation of MESP2-mCherry and UNCX-YFP signals in a Somitoid over time. Bottom, abscissa-position of the trough of the spatial auto-correlation function, indicating the typical cluster size, over time. **j**, Temporal plot (mean±95%CI) of mean squared displacement (n=3422 tracks from 2 Somitoids). **k-m**, Experiment scheme (k) and maximum-z-projection images (l, m) of re-aggregating MESP2/UNCX reporting Somitoids at 72h (1) or 96h (m). **n-p**, Experiment scheme (n) and maximum-z-projection images (o, p). After dissociation of 72h Somitoids, MESP2-high (top 10% mCherry) and MESP2-low (bottom 10% mCherry) single cells were separated by flow cytometry and re-aggregated. **q**, Illustration of AP polarity patterning in the Somitoid. Solid circles represent MESP2-high cells and hollow circles represent MESP2-low cells. Scale bars represent 50μm (h); 500μm (d, e, f); 500μm and 100μm in corresponding enlarged views (a, l, m, o, p).

How *MESP2* expression resolves from its initial wide segmental domain which marks the future somite to an anterior half-somite stripe defining the future anterior somite compartment is not understood. To see if our Somitoid system could help shed light on this process, we analyzed the dynamics of *MESP2* expression during AP patterning in vitro using the MESP2 reporter line (Extended Data Fig.2a). The temporal profile of the reporter suggests a rapid activation of *MESP2* from ∼64h to 72h (Fig. 2b). This phase is followed by a stabilization of the reporter expression in a salt-and-pepper pattern, spanning a 10-fold range of intensities (Fig.2f,g, Extended Data Fig.2f). These observations contrast with the established notion of an initial uniform *MESP2* expression in all cells of the future segmental domain^1,2^. Time lapse movies showed that, after 72h, cells progressively sorted together according to their *MESP2* expression levels defined by mCherry intensity. This led to the gradual formation of *MESP2*-high and *MESP2*-low clusters which eventually formed independent rosettes (Fig.2f,h, Extended Data Fig.2g, Supplementary Video5). To characterize this process, we measured the spatial auto-correlation of the mCherry (*MESP2* levels at 72h) and emerging YFP (*UNCX*) signals (Fig.2i, Extended Data Fig.2h,i). Before ∼80h, the auto-correlation functions were merely decreasing, suggesting the absence of a periodic spatial pattern (Fig.2i). At ∼80h, a trough formed at 90 microns and then quickly increased to 120 microns, suggesting a rapid formation of cell clusters. After ∼84h, the spatial auto-correlation function retained a damped oscillator-like shape, as typical for periodic patterns^20^. The onset of the periodic pattern that precedes rosette formation also corresponds to a slowing down of cell motility which can be visualized by measuring their mean squared displacement (Fig.2j). Thus, from 72h to 120h, the salt-and-pepper mCherry distribution became organized into mCherry-high and low clusters and then mCherry-high and low rosettes without further *MESP2* expression (Extended Data Fig.2j).

To test the role of cell sorting, we dissociated and re-aggregated Somitoids at 72h when cells are still mesenchymal, and at 96h when epithelial rosettes start emerging (Fig.2k, Extended Data Fig.2j). Rosettes mostly formed with either high mCherry or high YFP-expressing cells appeared in re-aggregates from 72h (Fig.2l), while homogeneous rosettes containing mixed YFP- and mCherry-positive cells were formed in re-aggregates from 96h (Fig.2m). Thus, cell sorting before epithelialization plays an important role in AP patterning of Somitoids. To investigate when AP fates in individual cells are determined in this process, we separated the *MESP2*-high or *MESP2*-low fractions from cultures dissociated at 72h and re-aggregated them separately (Fig.2n). At 120h, similar rosette morphogenesis was observed in both type of aggregates with *MESP2*-low re-aggregates expressing significantly higher level of *UNCX* than *MESP2*-high re-aggregates (Fig.2o,p, Extended Data Fig.2k). This suggests that AP cell fates are largely determined before cell sorting and rosette formation. Altogether, our experiments show that an initial heterogeneity of *MESP2* expression levels is translated into defined compartments of anterior and posterior identity via an active cell sorting mechanism (Fig.2q).

To further test whether such a mechanism explains the AP polarization of somites, we next set out to establish an in vitro model reproducing the spatial features of somitogenesis, including PSM elongation and sequential formation and patterning of somites. We treated iPSCs with CHIR and LDN for 24h, and then dissociated the cultures to single cells to generate spheroids using low adhesion wells (Fig.3a). We then embedded these spheroids into low-percentage Matrigel (10%) at 48h and cultured them in N2B27 media. By 96h, initially symmetric spheroids become elongated and develop into rod-shaped tissues exhibiting somite-like rosettes at one extremity (Fig.3b,c). Time lapse movies showed that these rosettes form sequentially starting from one end (which we define as anterior) while the other unsegmented end (the posterior end) kept extending (Extended Data Fig.3a,b, Supplementary Video6). The posterior end sometimes appeared bifurcated (Supplementary Video6). We termed these structures “Segmentoids”. Live imaging of a differentiating *PAX3-YFP* reporter line showed that *PAX3* expression initiated from the anterior end and propagated towards the posterior growing end accompanying rosette formation, indicating sequential maturation of the Segmentoids (Extended Data Fig.3c,d). TBXT/SOX2-positive cells were scattered in the spheroids at 48h (Fig.3d, Extended Data Fig.4). At 72h, TBXT/SOX2-positive cells congregated at the posterior end of the elongating Segmentoids, where they remained up to 96h. At 120h, we could barely detect TBXT while SOX2-only positive cells assembled into neural tube-like structures at the posterior tip of the tissue. These data suggest that the posterior growing end of the segmentoids resembles the tail bud end of embryos which contains the SOX2/TBXT positive Neuro-Mesodermal Progenitors (NMPs)^21^.

**Fig. 3.**
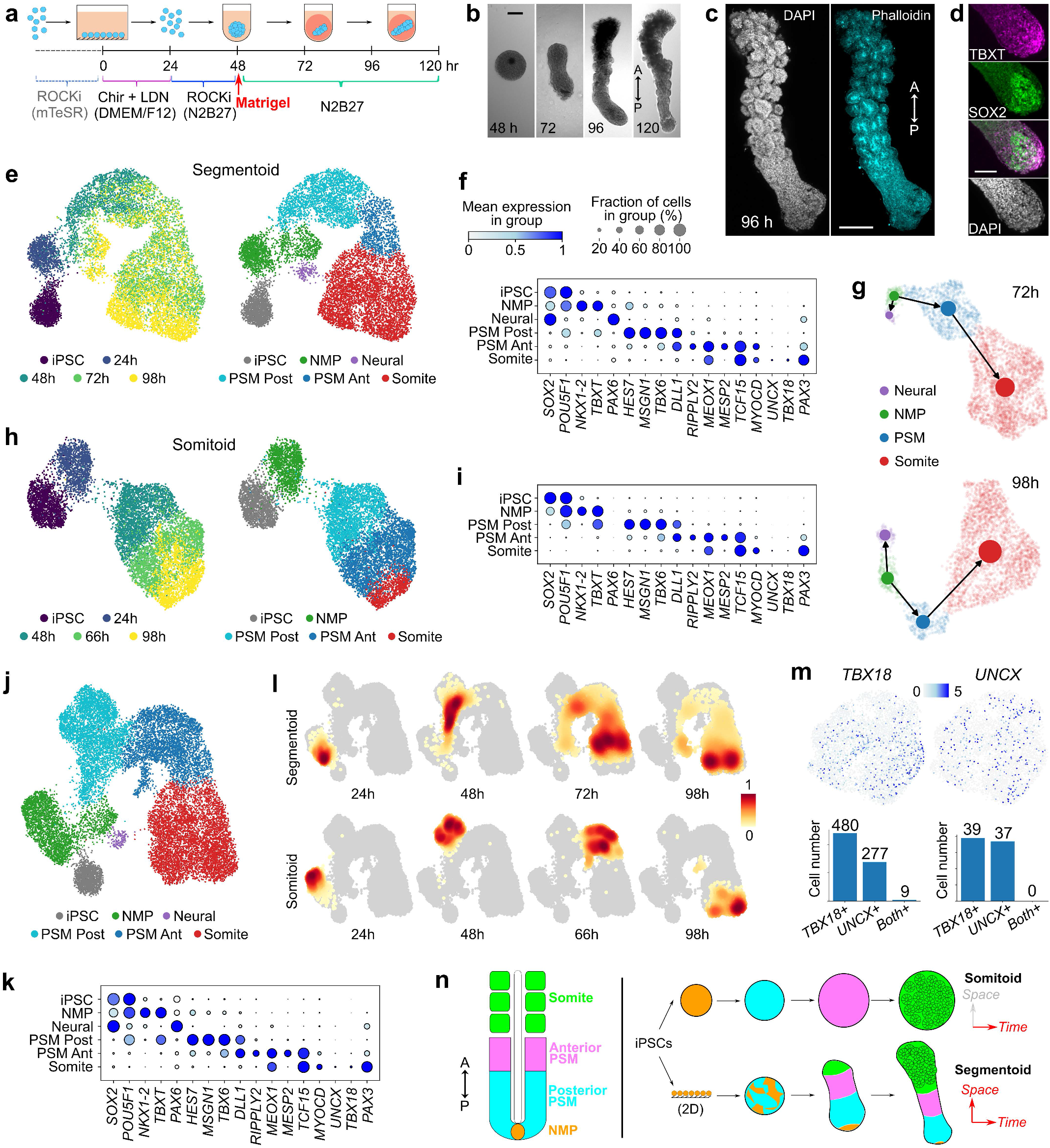
single-cell RNAseq characterization of the Segmentoid and Somitoid models. **a**, Illustration of the Segmentoid protocol. **b**, Representative bright field images of the Segmentoids at various time points. A, anterior; P, posterior. **c**, Maximum z-projection images of a 96h Segmentoid. **d**, Confocal images of the posterior tip of a 96h Segmentoid immunostained with TBXT and SOX2. **e**, UMAP embedding (10,861 cells) colored with Segmentoid timepoints (left) and cell types (right) identified with Leiden clustering. iPSC, 1491 cells; 24h, 1066 cells; 48h, 1577 cells from 76 Segmentoids; 72h, 3539 cells from 64 Segmentoids; 98h, 3188 cells from 32 Segmentoids. **f**, Dot plot of selected signature genes in cell type clusters from Segmentoids. The mean expression of each cluster is scaled per gene. **g**, PAGA graphs with velocity-directed edges in 72h (top) and 98h (bottom) Segmentoids. **h**, UMAP embedding (8,690 cells) colored with Somitoid timepoints (left) and cell types (right) following Leiden clustering. iPSC, 1491 cells; 24h, 1265 cells from 96 Somitoids; 48h, 2335 cells from 96 Somitoids; 66h, 2246 cells from 80 Somitoids; 98h, 1353 cells from 48 Somitoids. **i**, Dot plot of selected genes in cell type clusters from Somitoids. **j**, UMAP embedding of cells merged from both models (19,551 cells) colored with cell types identified with Leiden clustering. **k**, Dot plot of selected genes in cell type clusters from both models. **1**, Heatmap of cell density in UMAP embedding (scaled per timepoint). **m**, Top, somite sub-cluster highlighting cells expressing TBX18 (left) and UNCX (right); Bottom, number of cells expressing TBX18, UNCX, or both in Segmentoids (left) and Somitoids (right). **n**, comparison of the embryo with the two in vitro models. Each cell type is represented by the same color. Scale bars represent 200μm (b, c) and 100μm (d).

We next used single-cell RNA sequencing (scRNAseq) to characterize the identity and developmental trajectory of Segmentoids cells. Using the 10X Chromium v3.1 platform, we sequenced a total of ∼10,000 cells including iPSCs and Segmentoids at 24h, 48h, 72h, and 98h. When all time points were merged and analyzed together, cells on the UMAP spontaneously organized into a developmental trajectory reflecting the progression of somitogenesis (Fig.3e,f). Cells were clustered using the Leiden algorithm and the identity of clusters was defined based on differentially expressed genes. The clusters included iPSCs, NMPs (expressing *SOX2, TBXT*, and *NKX1*.*2*), Posterior PSM (expressing *MSGN1, TBX6*, and *HES7*), Anterior PSM (expressing *TCF15* and *MESP2*), and Somite (expressing *PAX3, UNCX*, and *TBX18*). A small Neural cluster (expressing *SOX2* and *PAX6*) was also observed. NMP and PSM populations gradually decreased with time while the somite population increased (Extended Data Fig.5a). Velocity combined with PAGA analysis confirmed that both Neural and Mesodermal cells arise from the NMP progenitors (Fig.3g, Extended Data Fig.5b,c). We also observed a collinear expression pattern of *HOX* genes which terminated at the level of the *HOX9* group by 98h (Extended Data Fig.6a). Altogether, we have established a 3D system in which differentiating human iPSC recapitulate the spatiotemporal progression of somitogenesis.

We also performed a similar scRNAseq analysis of the Somitoids. We sequenced a total of ∼10,000 cells at 24h, 48h, 66h and 98h, and observed clusters similar to those of Segmentoids except for the neural cluster which was absent (Fig.3h,i). The activation of HOX gene clusters followed a similar temporal progression (Extended Data Fig.6b). In contrast to Segmentoids, cells from a defined time point could be ascribed to a single cluster (Fig.3h), indicating synchronized differentiation across the entire culture. We created a merged dataset containing all cells from the two systems. Cells from the two datasets with the same identity merged into one single cluster (Fig.3j,k), indicating that the cell types generated in the two systems are similar. Using density plots we showed that Somitoids time-points are clearly defined by a homogenous cell identity (Fig.3l). In contrast, the Segmentoid time points contain multiple differentiation stages (Fig.3l), recapitulating the progression of differentiation observed during somitogenesis in embryos. We extracted the somite population from the merged dataset and investigated the onset of the anterior and posterior identities focusing on the expression of *TBX18* and *UNCX* (Fig.3m). We found that the expression of the two genes occurred at the somite stage and was mutually exclusive (0/76 in Somitoids and 9/757 in Segmentoids were double positive cells). Yet, these cells did not segregate into distinct clusters suggesting that they share a similar transcriptome at these stages despite their different AP identities. These analyses demonstrate that both systems can recapitulate somitogenesis in vitro with Somitoids showing synchronized cell differentiation while Segmentoids exhibit a spatially organized progressive maturation similar to that of the embryonic tissue (Fig.3n).

We next investigated somite formation and patterning in Segmentoids, using a reporter cell line monitoring the expression dynamics of *HES7* (destabilized YFP), *MESP2* (mCherry) and *UNCX* (YFP) (Fig.4a-d, Extended Data Fig.7a). Oscillatory expression of *HES7* occurred in the posterior PSM and became down-regulated where *MESP2* expression started (Fig.4a,b, Supplementary Video7). The domain of *MESP2* expression progressed posteriorly in a staggered manner, closely in sync with *HES7* oscillations (Fig.4a,b). Time auto-correlation analysis shows oscillations with a period of 4.6±0.1 h for *HES7* and 5.4±0.5 h for *MESP2* (Fig.4d, Extended Data Fig.7b). Thus, the coupling between the segmentation clock and *MESP2* induction observed in mouse embryos^22^ is recapitulated in Segmentoids. At 120h, alternating stripes of mCherry (*MESP2*) and YFP (*UNCX*) were observed (Fig.4c). From posterior to anterior, mCherry first appeared as a broad stripe followed by narrower bands with complementary YFP bands emerging (Fig. 4c), recapitulating the expression patterns observed in mouse in vivo. Patterning was independent of morphogenesis since mCherry/YFP stripes were established in the presence of ROCKi which blocked rosette formation (Extended Data Fig.7c).

**Fig. 4.**
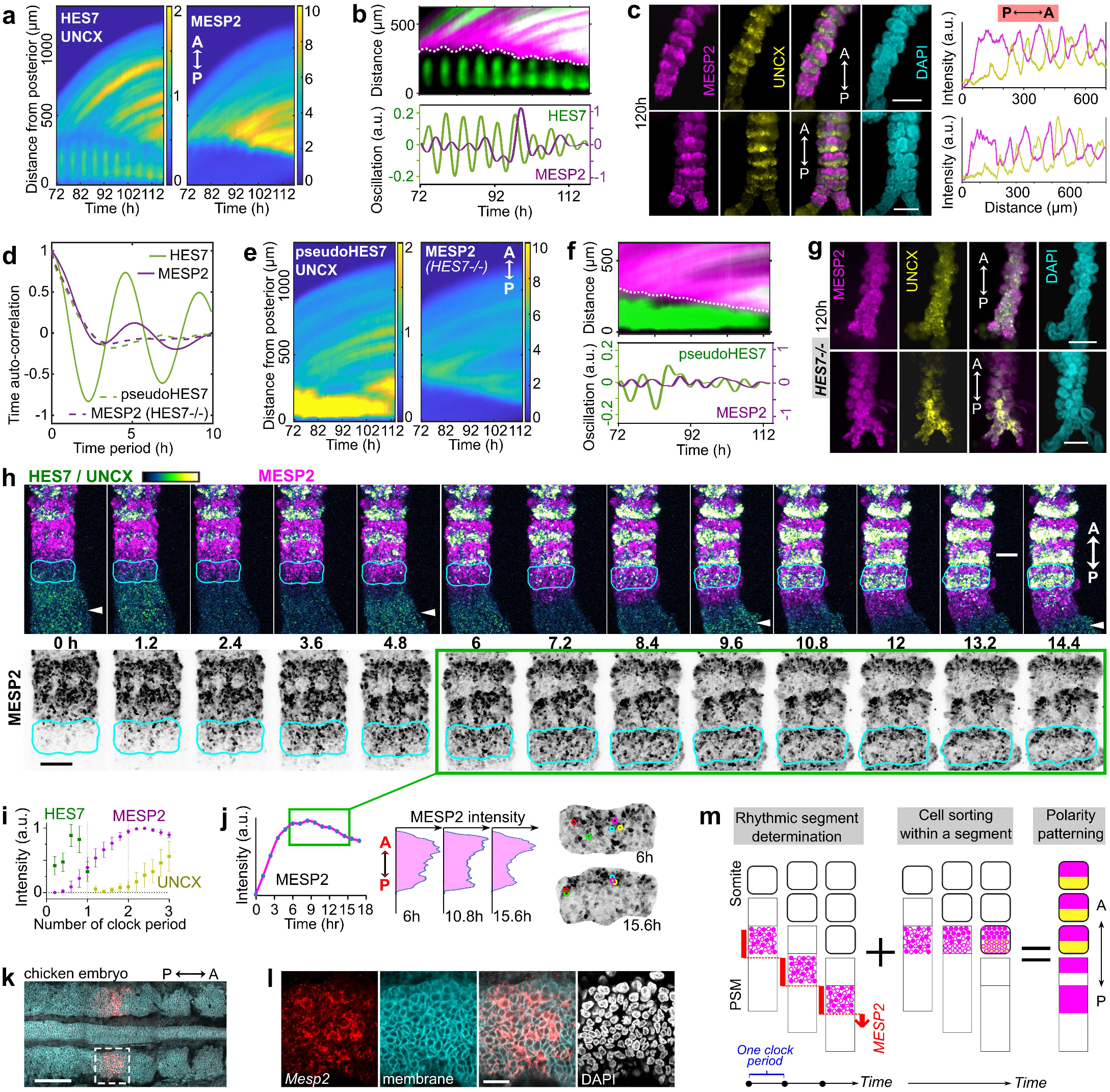
Formation of anterior and posterior somite compartments in Segmentoids. **a**, Kymographs of reporters for HES7 (posterior part in the kymograph), UNCX, and MESP2 in the same Segmentoid. Segmentoids are aligned to the posterior tip at each time point. **b**, Top, merged kymographs of HES7/UNCX (green) and MESP2 (magenta). Dotted line highlights the start of the MESP2 signal region. Bottom, HES7 and MESP2 oscillations (Methods). **c**, Wide-field images (left) of 120h MESP2/UNCX -reporting Segmentoids and intensity profiles (right) from posterior **(P)** to anterior (A) end along each Segmentoid. **d**, Average time auto-correlation of HES7 and MESP2 reporter oscillations in WT (n=7 Segmentoids) and HES7-null (n=6 Segmentoids). The first peak of the auto-correlation function (time>0h) indicates the oscillation period. **e**, Kymographs of reporters for pseudoHES7, UNCX, and MESP2 in the same HES7-null Segmentoid. **f**, Top, merged kymographs of pseudoHES7/UNCX (green) and MESP2 (magenta) in a HES7-null Segmentoid. Dotted line highlights the start of the MESP2 signal region. Bottom, pseudoHES7 and MESP2 oscillations (Methods). **g**, Wide-field images of 120h MESP2/UNCX-reporting HES7-null Segmentoids. **h**, Time-lapse, maximum-z-projection confocal images of MESP2 and UNCX/HES7 reporters (top) and enlarged grey-scale images of MESP2 (bottom) in a Segmentoid. Cyan solid outlines indicate the same forming segment and white arrowheads indicate the approximate peaks of HES7 oscillation. **i**, Reporter dynamics (mean±s.d.) in forming segments aligned according to phases of HES7 oscillation (n=6 segments in 2 Segmentoids). **j**, Left, temporal profile of MESP2 intensity in the forming segment outlined in h. Green solid-line boxes indicate the corresponding time points. Middle, MESP2 profiles along a line scan of the AP axis of the same segment. Right, isolated segments with circles of the same color indicating the same cells. **k**, Merged confocal image of a chicken embryo stained with MESP2 HCR probe (red) and a membrane dye (cyan). **l**, Enlarged view of the region indicated by the dotted-line box ink. **m**, Model illustration: The stepwise, salt-and-pepper induction of MESP2 expression organized by the segmentation clock defines a maturing region at the anterior PSM; Cell sorting within this forming segment rearranges MESP2-high cells to the anterior compartment while MESP2-low cells to the posterior, which further express differential genes to achieve AP polarity patterning. Scale bars represent 200μm (c, g), 100μm (h, k) and 20μm (1).

To investigate the role of the segmentation clock in somite polarity patterning^23^, we examined *HES7*-null Segmentoids. The YFP signal, reporting activity of the *HES7* promoter in absence of HES7 protein (pseudoHES7), was confined to the posterior tip of the elongating tissue. It progressively shrank in a non-oscillatory pattern as the end grew (Fig.4e,f, Supplementary Video8). The onset of *MESP2* expression was still coordinated with the arrest of pseudoHES7 in space. In contrast to its staggered progression in WT Segmentoids, the *MESP2* expression domain moved continuously towards the posterior end in the *HES7*-null mutants (Fig.4d,f). At 120h, no alternating stripes of mCherry (*MESP2*) or YFP (*UNCX*) could be observed in the mutant (Fig.4g). Cells of posterior and anterior identity appeared randomly distributed with some clusters formed, consistent with the segmental polarity defects reported in *HES7*-null mouse embryos^24^. To confirm this apparent disorganization in *HES7*-null Segmentoids, we used the nematic order parameter^25^ of the *MESP2/UNCX* signal as a measure of anisotropy; we found that the nematic order parameter was lower in *HES7*-null Segmentoids than control during differentiation (Extended Data Fig. 7d). Thus, as with Somitoids, the segmentation clock is not required for the expression of AP identity genes in individual cells, but its output conferring rhythmicity to *MESP2* induction and segment determination appears to play an important role in the spatial organization of stripes of anterior and posterior identity in the forming somites.

To explore whether the cell sorting mechanism observed in the Somitoids could be involved in the generation of alternate AP stripes, we analyzed the formation of individual segments in Segmentoids. We first observed a segment-wide transient expression of *MESP2* lasting for ∼1 clock period followed by *UNCX* expression in *MESP2*-low cells during the next clock period (Fig.4h,i, Supplementary Video9). As with Somitoids, the initial induction of *MESP2* resulted in cells displaying a broad distribution of expression levels throughout the newly specified segment (Fig.4h). Cells of high *MESP2* expression levels gradually congregated to the anterior compartment while the overall mCherry intensity in the segment stayed constant (Fig.4j, Extended Data Fig.7e, Supplementary Video10), suggesting that no new *MESP2* expression occurred during segregation of the anterior and posterior domains. Spatial auto-correlation analysis following the same segment showed an emerging pattern of mCherry during this time window (Extended Data Fig.7f). Thus, this suggests that formation of the stripes of the anterior and posterior somitic compartments, does not rely on differential regulation of *MESP2* expression as usually inferred. To test whether such a heterogeneous *MESP2* expression is observed in vivo, we used the quantitative *in situ* Hybridization Chain Reaction (HCR), to examine the onset of *MESP2* expression in the anterior PSM of chicken and mouse embryos. Indeed, we observed a clear salt-and-pepper pattern of *MESP2* expression among cells of the future segmental domain indicating that the sorting mechanism that we uncovered in vitro is likely operating in vivo (Fig.4k,l, Extended Data Fig.7g,h).

In summary, we established two iPSC-derived 3D models recapitulating human somitogenesis, Somitoids and Segmentoids (Fig.3n). In contrast to gastruloids or Trunk-Like Structures^17,26,27^ which harbor cell lineages derived from the three germ layers, our two models contain almost exclusively paraxial mesoderm. Somitoids recapitulate the temporal sequence of somitogenesis, with all cells undergoing differentiation and morphogenesis in a synchronous manner. This system can provide unlimited amounts of cells precisely synchronized in their differentiation. It will allow exploring these patterning processes at an unprecedented level of detail. On the other hand, Segmentoids reconstruct the spatio-temporal features of somitogenesis, including gene expression dynamics, tissue elongation, sequential somite morphogenesis, and polarity patterning. They therefore provide an excellent proxy to study human somitogenesis.

Together our work suggests a novel framework (Fig.4m) explaining how somite AP polarity is coordinated with segmental determination. We show that acquisition of somite AP identities in individual cells is established early in the nascent segmental domain in a salt-and-pepper fashion and does not require the segmentation clock, tissue elongation, or somite epithelialization in vitro. Our data suggest that two sequential processes are required for establishing somite AP polarity (Fig.4m): 1-the staggered initiation of *MESP2* expression in a salt and pepper fashion in defined segmental stripes specified by the segmentation clock; 2-the sorting of cells according to their *MESP2* expression levels to form the AP compartments of the future somite. This sorting mechanism identified in vitro appears to be conserved in vivo. The patterning mechanism ensuring that the *MESP2*-high compartment is anterior rather than posterior remains to be identified but could rely on the gradients of FGF/Wnt and RA along the PSM. Thus, our work exemplifies how the resolution offered by PSC-derived in vitro systems can be used to answer long-standing developmental biology questions.

## Acknowledgements

We thank Sudhir Gopal Tattikota from N. Perrimon lab for help with scRNA sequencing experiments. We thank the Biopolymers Facility at Harvard Medical School for providing 10X Genomics Chromium Controller instrument access and sequencing consultation. We thank the NeuroTechnology Studio at Brigham and Women’s Hospital for providing microscope access and consultation on data acquisition and data analysis. We thank the Harvard Neurobiology Imaging Facility for access to the FV1000 confocal microscope (NINDS P30 Core Center grant NS072030). We thank S. Megason for critical reading of the manuscript. Research in the Pourquié lab was funded by a grant from the National Institute of Health (5R01HD085121). Y.D. is supported by Fondation pour la Recherche Médicale (FRM) PLP2020100012456.

## Author contributions

Y.M. designed, performed, and analyzed most biological experiments; Y.D. analyzed scRNA seq data; A.D. developed the codes and performed quantitative image analysis with S.D.; K.Z. performed RNA seq sample preparation and data analysis; A.S. and J.G.L. conducted embryo HCR experiments with help from L.S.; J.R. and O.A.T. contributed to scRNA experiments; A.S., A.M., P.R., and M.D.-C. contributed to data analysis or experiments. Y.M., Y.D., and O.P. wrote the manuscript with inputs from all authors; and O.P. supervised the study.

---

**Extended Data Fig.1.**
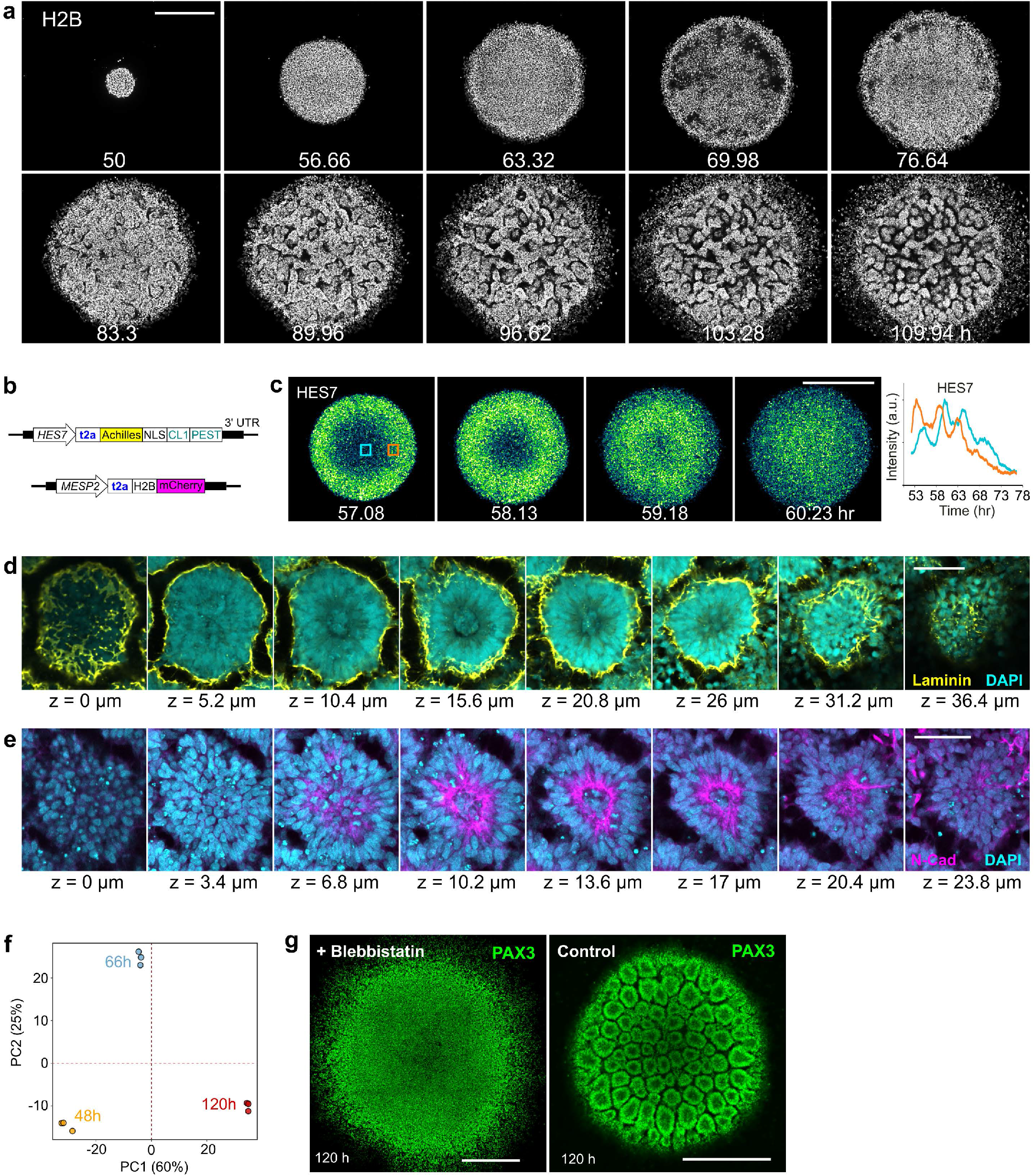
Characterization of Somitoids. **a**, Time-lapse confocal images of H2B-mCherry in a spreading Somitoid. **b**, Illustration of the design of the HES7/MESP2 double-reporter cell line. **c**, Left, time-lapse confocal images of HES7 wave; Right, temporal profiles of HES7 reporter in two different regions indicated by the blue and orange boxes. **d, e**, Confocal slices from the bottom (z=0 μm) to the top of a rosette in 120h Somitoid stained with Laminin (d) and N-Cadherin (e). **f**, Principal components analysis using the same RNA sequencing datasets shown in Fig. 1h. **g**, Confocal images of 120h PAX3-reporting Somitoids treated with 5μM Blebbistatin (left) and control (right). Scale bars represent 500μm (a, c, g) and 50μm (d, e).

**Extended Data Fig.2.**
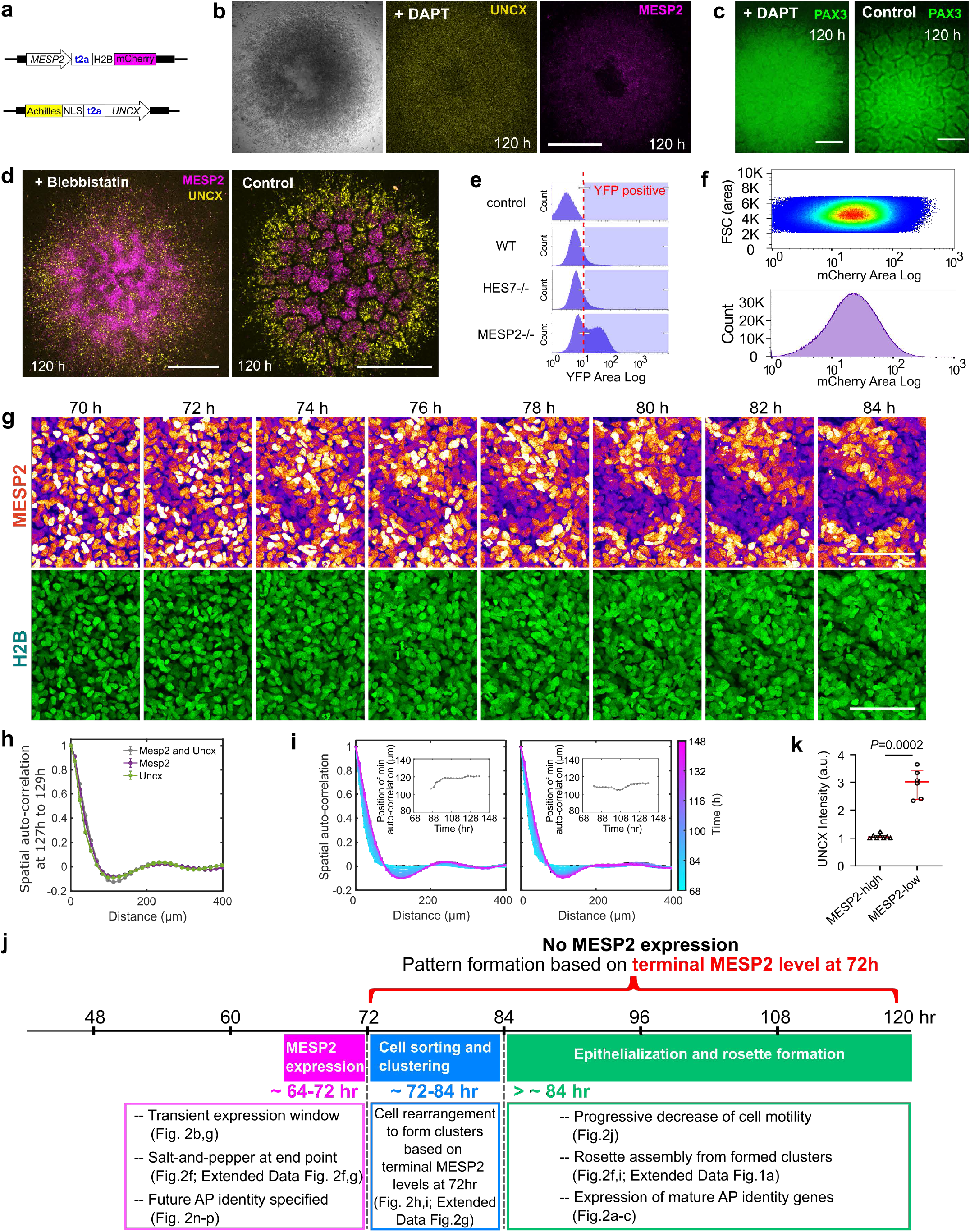
Antero-Posterior patterning in Somitoids. **a**, Illustration of the design of the MESP2 and UNCX reporters, **b**, Images of an UNCX and MESP2 reporting Somitoid treated with 50μM DAPT. **c**, Wide-field images of PAX3-reporting Somitoids treated with 50μM DAPT (left) and control (right), **d**, Maximum-z-projection confocal images of UNCX and MESP2 reporting Somitoids treated with 5μM Blebbistatin (left) and control (right), **e**, Histograms of flow cytometry analysis of UNCX-YFP in 120h Somitoids (control, WT, HES7-null, and MESP2-null cell lines) with debris and doublets removed. Control is the parental NCRM1 cell line. Fractions on the right side of the red dotted line in the histograms are defined as YFP-positive. **f**, Scattered plot (top) and histogram (bottom) of flow cytometry analysis on MESP2-mCherry Somitoids at 72h with debris and doublets removed, **g**, Time-lapse maximum-z-projection confocal images of MESP2 reporter (top) and H2B-GFP (bottom) in the same region of a Somitoid. **h**, Spatial auto-correlation (sole MESP2 signal, sole UNCX signal or them combined together) once rosettes are formed (representative example from n=3 Somitoids). **i**, Additional examples of spatial auto-correlation analysis and abscissa-position of the auto-correlation trough (inset) of MESP2/UNCX double reporting Somitoid over time, **j**, Summary of MESP2 expression and pattern formation processes in the timeline of the Somitoid differentiation, **k**, Quantification of UNCX reporter in MESP2-high (n=8 re-aggregates from 3 experiments) and MESP2-low (n=6 re-aggregates from 3 experiments) re-aggregates in Fig.2n-p, paired two-tailed t-test. Scale bars represent 500μm (b, d), 200μm (c), and 100μm (g).

**Extended Data Fig.3.**
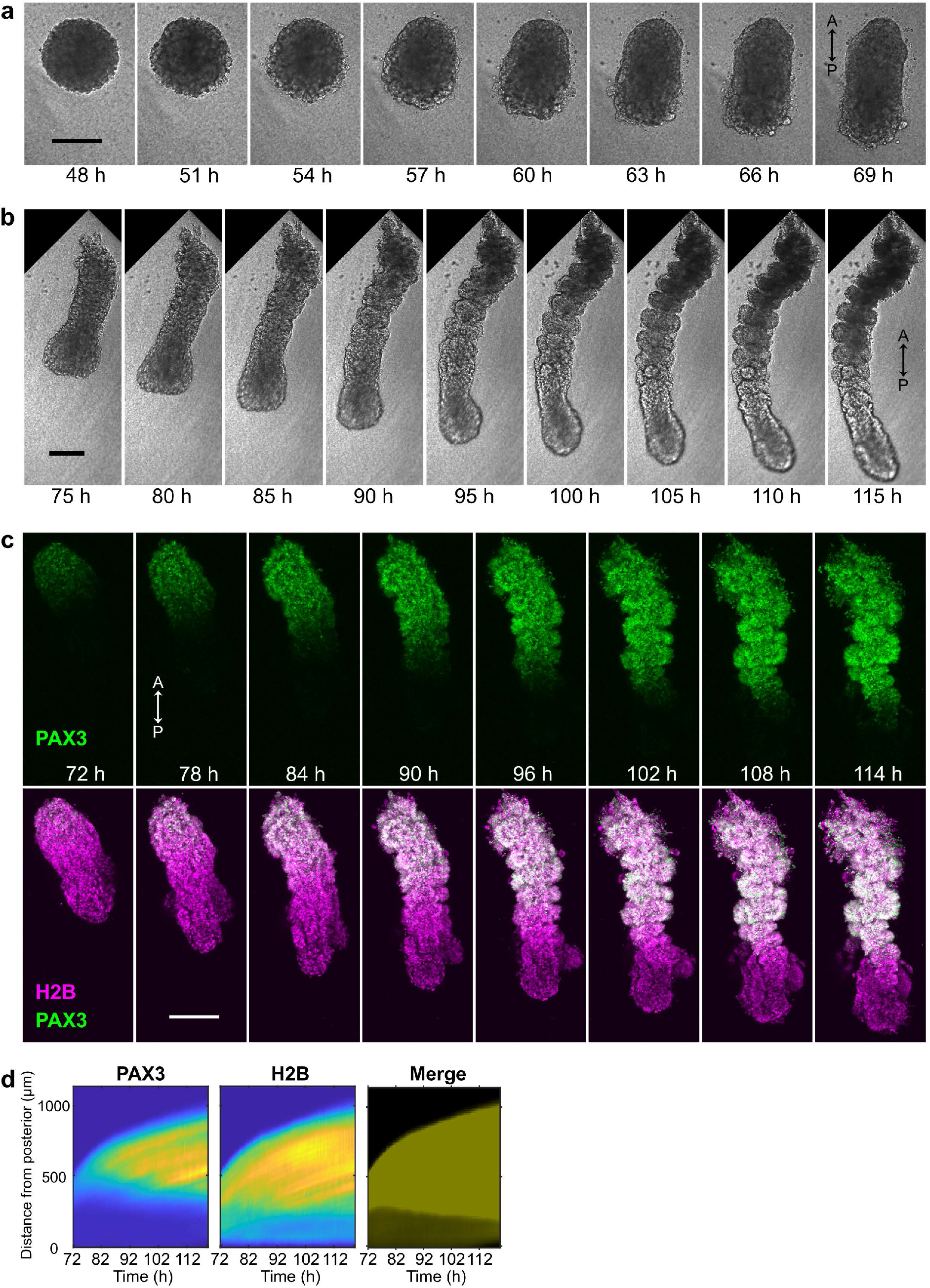
The Segmentoid model. **a, b**, Time-lapse bright field images of the Segmentoid model. A, anterior; P, posterior. **c**, Time-lapse maximum-z-projection confocal images of PAX3-YFP reporter (top) and PAX3-YFP merged with H2B-mCherry (bottom) in a Segmentoid. **d**, Kymographs of PAX3 reporter (left), H2B (middle), and merged channels (right) in the same Segmentoid. Segmentoids are aligned to the posterior tip at each time point. All scale bars represent 200μm.

**Extended Data Fig.4.**
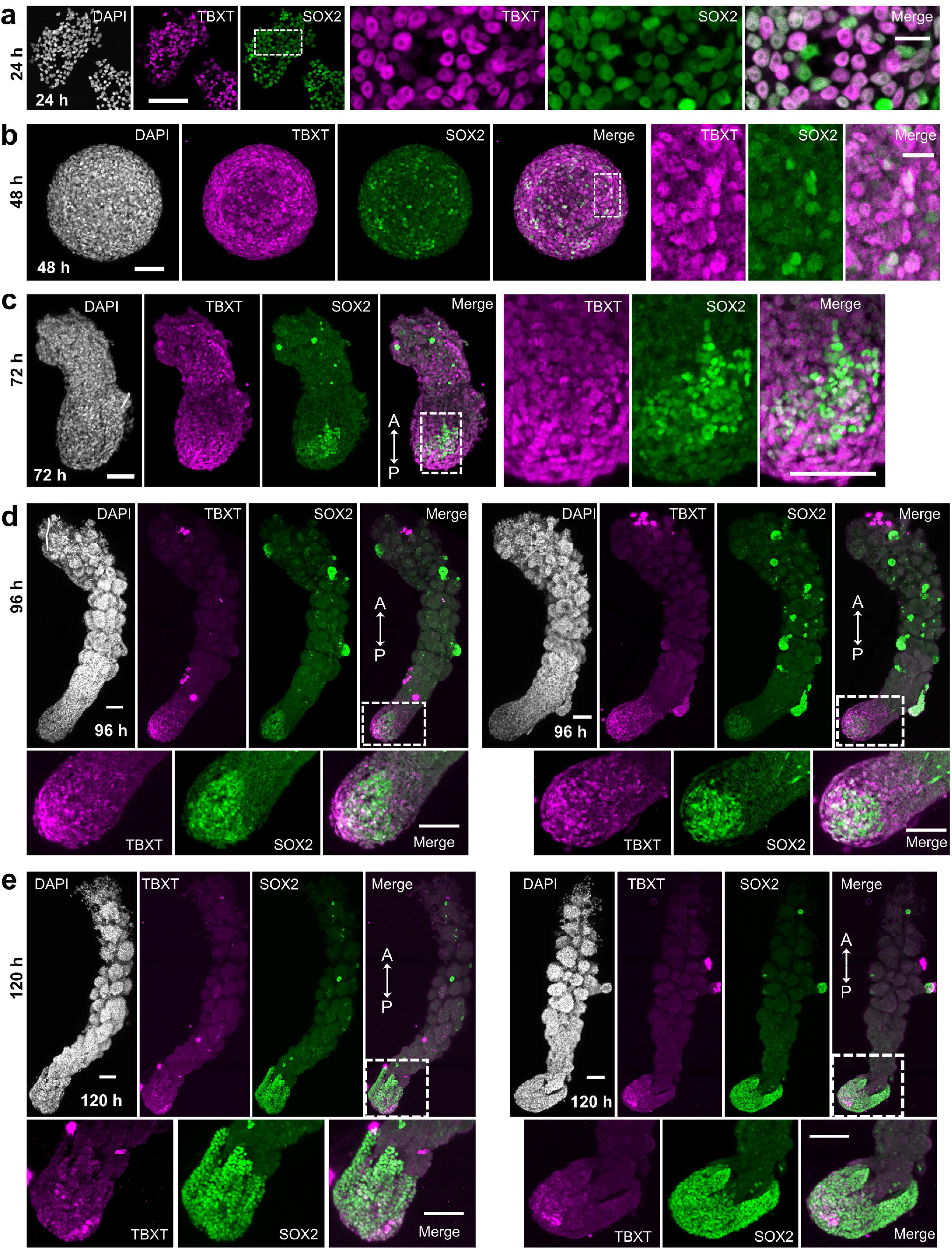
Expression of TBXT and SOX2 in Segmentoids. Confocal images of immunostaining of TBXT and SOX2 at 24h (**a**), 48h (**b**), 72h (**c**), 96h (**d**), and 120h (**e**) of the Segmentoid model. Maximum-z-projection images are shown from b-e. A, anterior; P, posterior. Scale bars (a, b) represent 100μm and 20μm in corresponding enlarged views; Scale bars (c, d, e) represent 100μm.

**Extended Data Fig.5.**
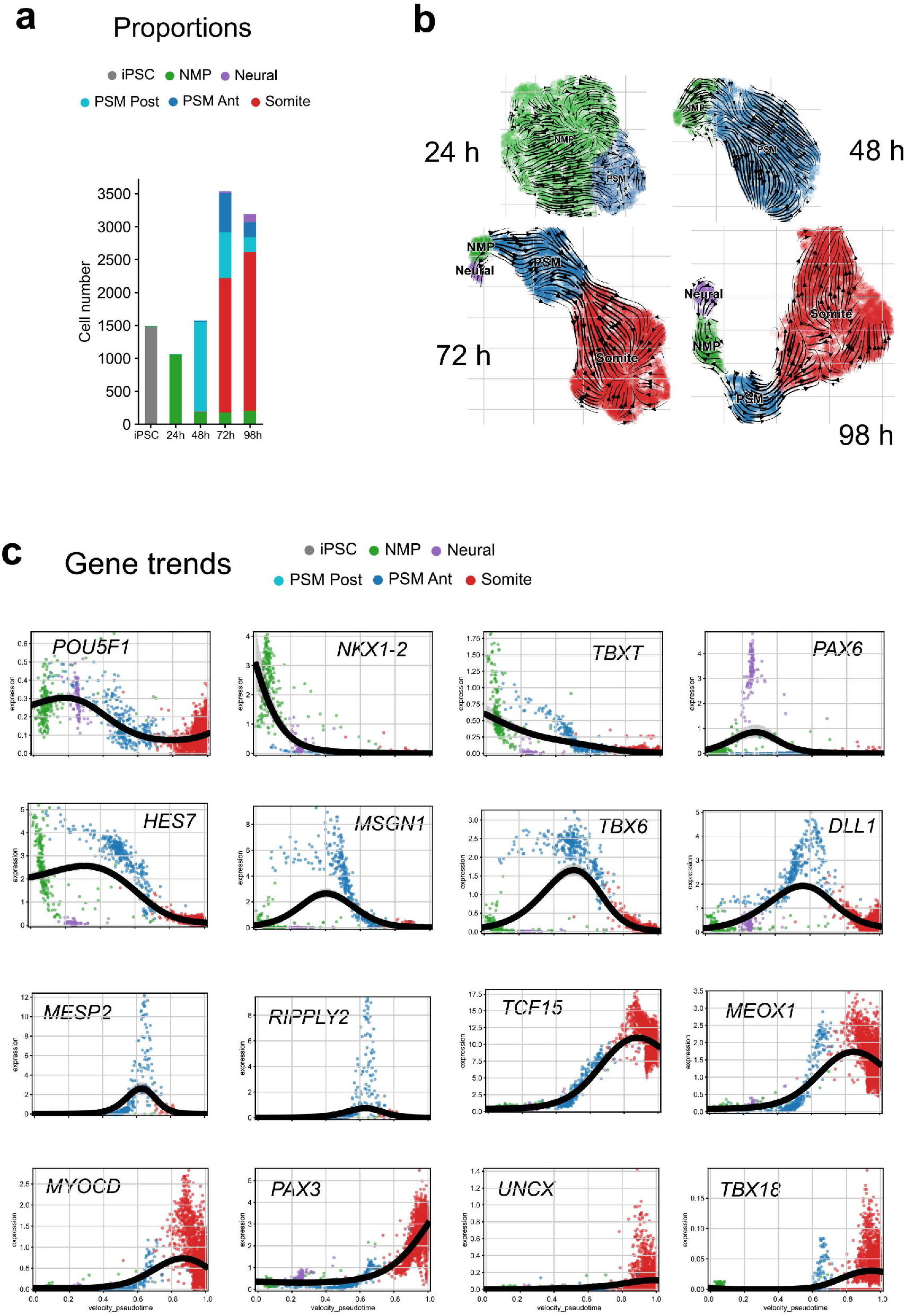
Single-cell RNAseq of the Segmentoid model. **a**, Proportion of cell types identified with Leiden clustering at different timepoints of the Segmentoid model. **b**, Stream plots of velocities on the UMAP after correction for differential kinetics recapitulating trajectory of cell types at various timepoints. **c**, Signature gene expression trends (Log2/Normalized) toward somite as the specific terminal population.

**Extended Data Fig.6.**
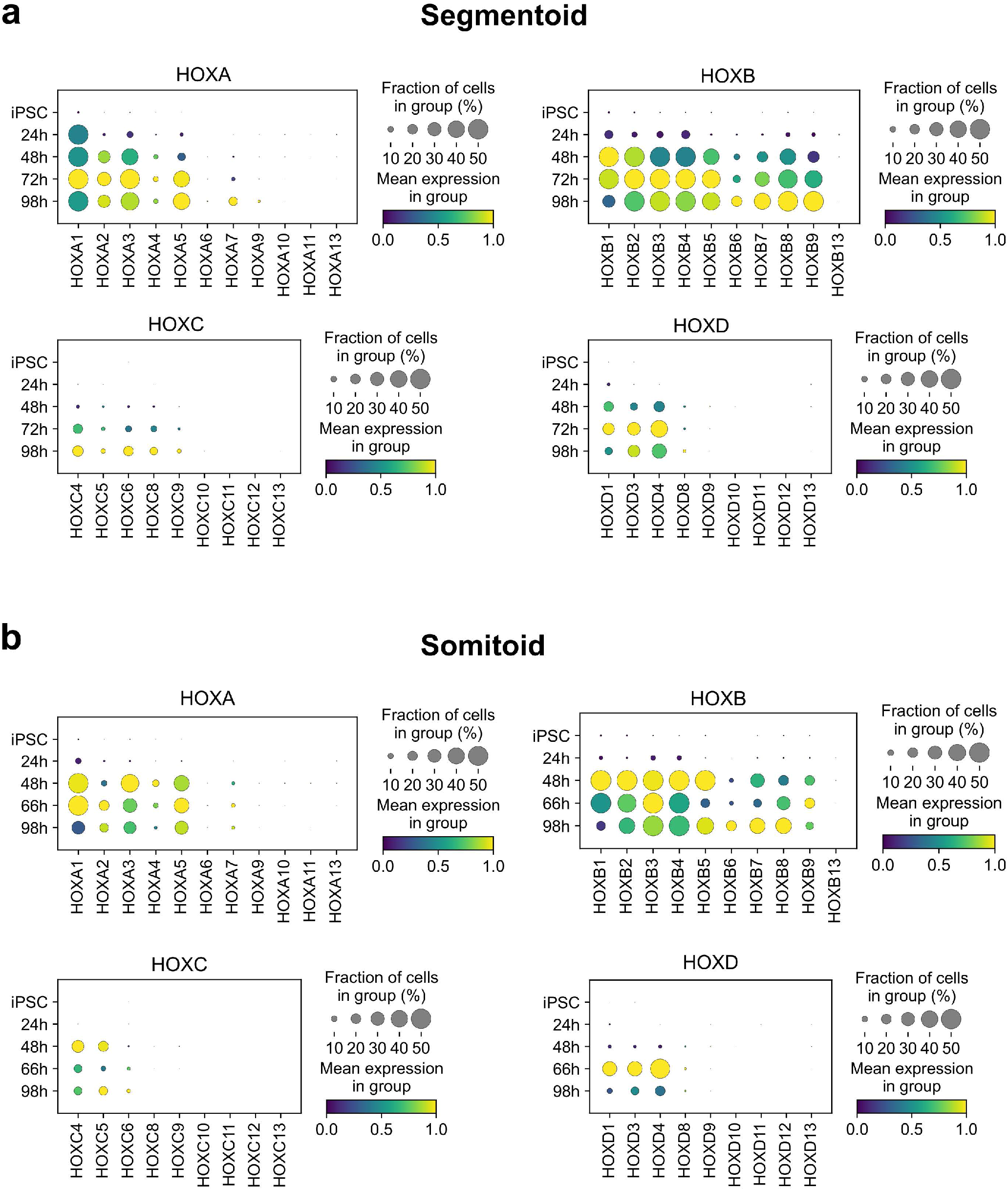
HOX genes expression in the in vitro models. Dot plots of HOX-family genes expression at various timepoints of the Segmentoid model (**a**) and the Somitoid model (**b**). The mean expression of each cluster is scaled per gene.

**Extended Data Fig.7.**
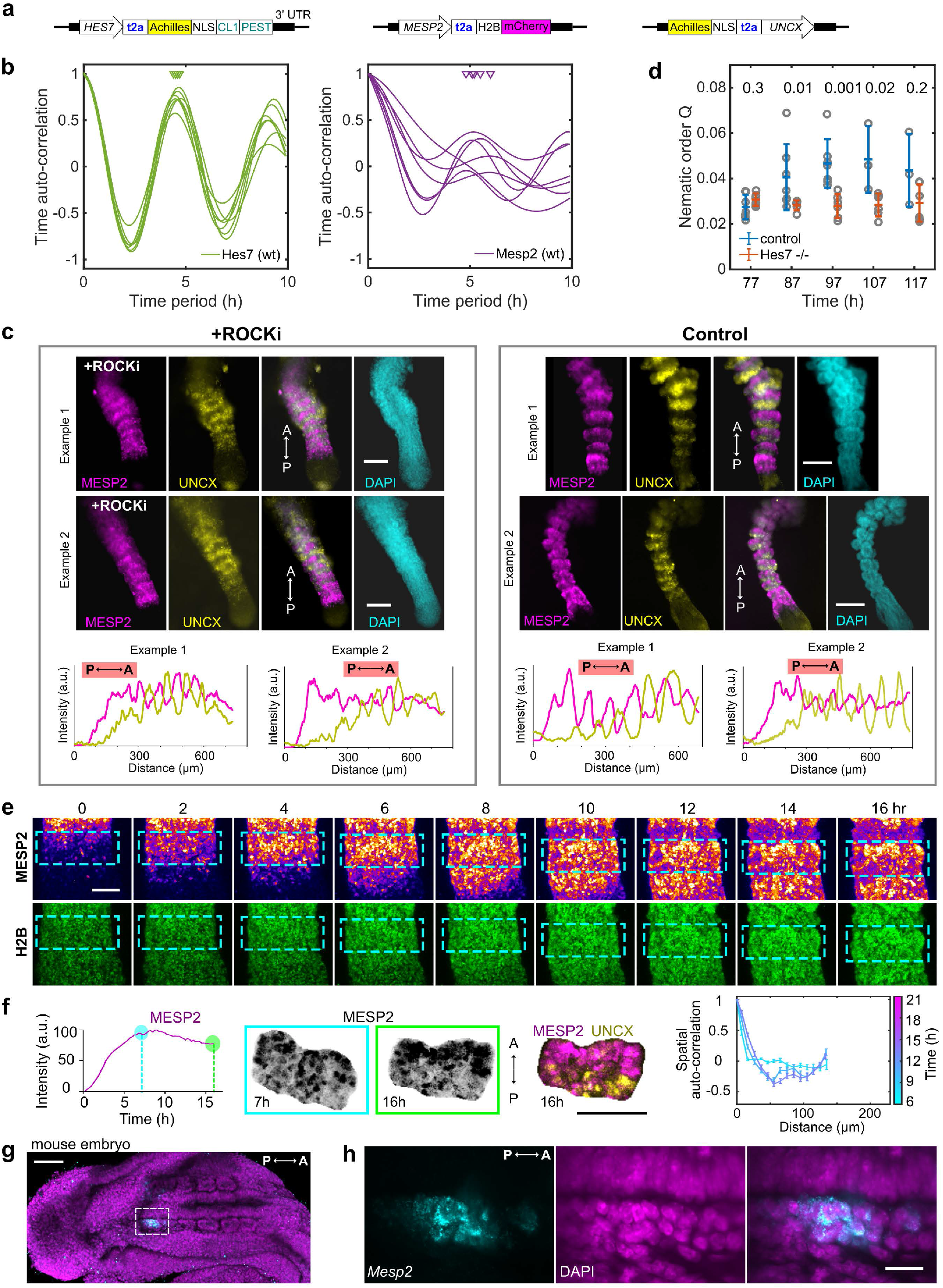
Antero-Posterior patterning in Segmentoids. **a**, Illustration of the design of the triple-reporter cell line. **b**, Time auto-correlation of HES7 and MESP2 reporter oscillations in individual WT Segmentoids (see Fig. 4d). Triangles indicate auto-correlation peaks, which in tum indicate oscillation period. **c**, Wide field images and graphs of reporter intensities from posterior (P) to anterior (A) end along 120h Segmentoids treated with 10μM ROCKi (left) and controls (right). **d**, Average nematic order of MESP2/UNCX signals in WT and HES7-null Segmentoids as a function of time. Statistics was performed with a Wilcoxon rank-sum test and P-value is shown. **e**, Time-lapse maximum-z-projection confocal images of MESP2 reporter (top) and H2B-GFP (bottom) in a Segmentoid. Cyan dotted-line boxes indicate the same developing segment. **f**, Temporal profile of MESP2 intensity (left), maximum-z-projection confocal images (middle), and spatial auto-correlation analysis of MESP2 and UNCX reporters as a function of time (right) in an individual developing segment in a Segmentoid. **g**, Merged maximum-z-projection confocal image of a mouse embryo stained with Mesp2 HCR probe (cyan) and DAPI (magenta). **h**, Enlarged view of the region indicated by the dotted-line boxing. Scale bars represent 200μm (c), 100μm (e, f, g) and 20μm (h).

